## Supplementary File 1 for "Genetic integration of behavioural and endocrine components of the stress response"

**Supplementary File 1: Selection of OFT behaviours**

Behavioural traits from the Open Field Trials (OFTs) were selected based upon previous research in this population and typical OFT measurements that describe variation in movement behaviour. Here we briefly describe selection of some traits against potential alternatives.

i. ‘Freezing’ behaviour.

Freezing behaviour is important to the ‘coping styles’ model as this is often used to describe the reactive style of coping. A velocity threshold for active swimming (4cm/s) has been defined for this population and used in previous studies for percentage of time spent active and for the number of freezings (e.g., White & Wilson 2018). We assessed ‘inactivity’ (i.e., 100 – time spent active, the percentage of time spent below the velocity threshold), but this was strongly correlated with ‘track length’ at the observation level (*r* = -0.96, 95% CI = [-0.97,-0.96], t = -210.3, *P* < 0.001; Figure 1) and so was considered not to add any further value. We also fit a bivariate animal model (with fixed and random effects as in the main text), and found that all correlations were >0.86, including a genetic correlation of *r*_A_ = 0.99 ± 0.01. We elected to retain track length rather than inactivity because it is a raw value rather than percentage, and so provides useful information on (for example) extreme values of track length.

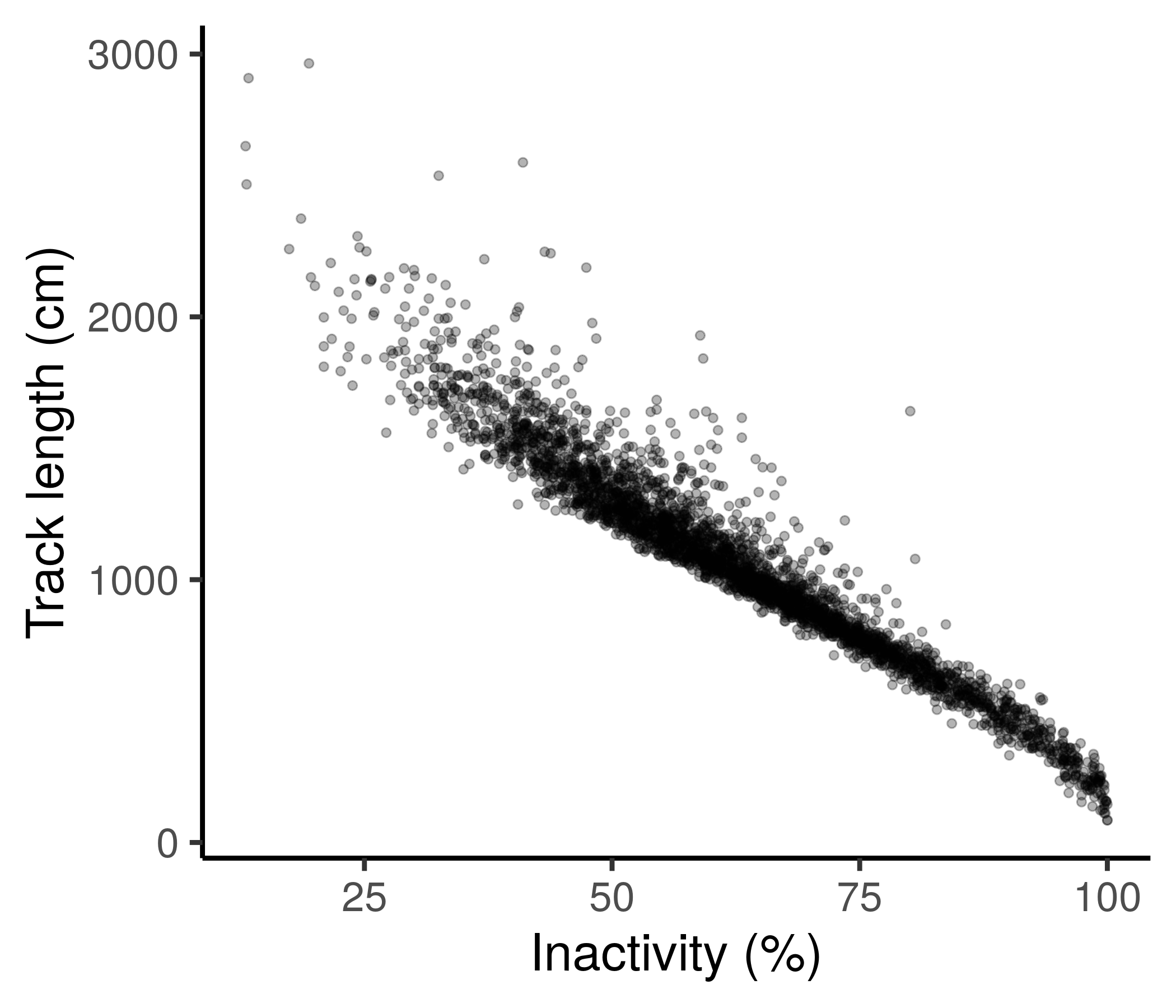

Figure 1: Percentage of time inactive is strongly correlated with track length at the phenotypic level.

ii. Thigmotaxis

OFTs are often used for research in personality traits (such as ‘boldness’ or ‘exploration’) as well as for anxiety behaviours. In the former, it is typical to define a central zone and quantify the time spent exploring this region as a measure of boldness. In the latter, it is typical to consider the average distance from the arena wall. Here we found that these measurements are highly correlated at the observation level (*r* = 0.94, 95% CI = [0.94,0.94], t = 159.2, *P* < 0.001; Figure 2). We also fit a bivariate animal model (with fixed and random effects as in the main text), and found that all correlations were >0.9, including a genetic correlation of *r*_A_ = 0.99 ± 0.01. These results suggested no gain to interpretation of using both, and we elected to retain time in the middle as the zoning approach is standard in much animal personality work.

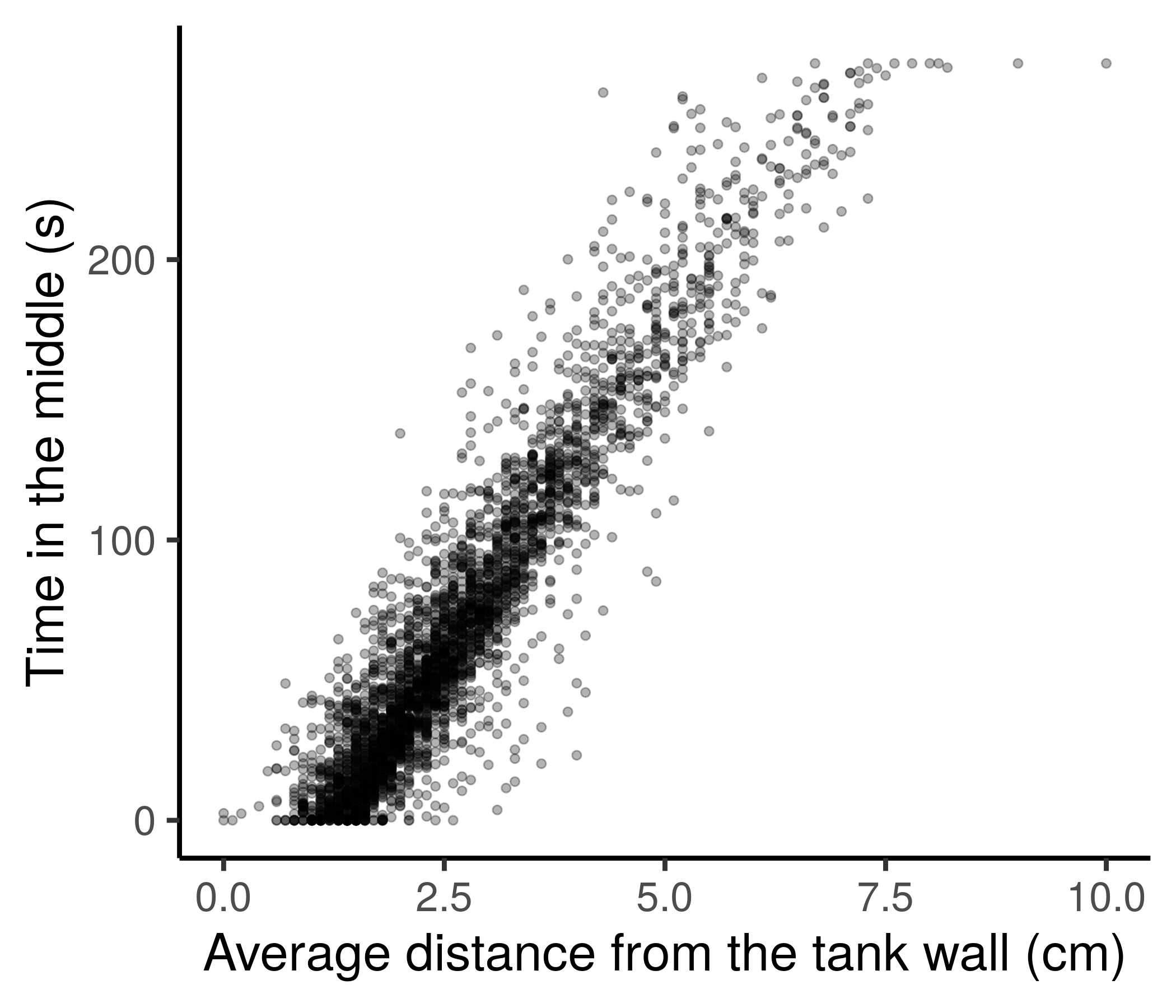

Figure 2: Time in the middle is strongly correlated with average distance from the tank wall at the phenotypic level.
